# Transcriptional, epigenetic, and functional reprogramming of blood monocytes in non-human primates following chronic alcohol drinking

**DOI:** 10.1101/2021.05.12.443856

**Authors:** Sloan A. Lewis, Suhas Sureshchandra, Brianna Doratt, Vanessa A. Jimenez, Cara Stull, Kathleen A. Grant, Ilhem Messaoudi

**Author notes:** Corresponding Author: Ilhem Messaoudi, Molecular Biology and Biochemistry, University of California Irvine, 2400 Biological Sciences III, Irvine, CA 92697.

## Abstract

Chronic heavy drinking (CHD) of alcohol is a known risk factor for increased susceptibility to bacterial and viral infection as well as impaired wound healing. Evidence suggests that these defects are mediated by a dysregulated inflammatory response originating from myeloid cells, notably monocytes and macrophages, but the mechanisms remain poorly understood. Our ability to study CHD is impacted by the complexities of human drinking patterns and behavior as well as comorbidities and confounding risk factors for patients with alcohol use disorders. To overcome these challenges, we utilize a translational rhesus macaque model of voluntary ethanol self-administration that closely recapitulates human drinking patterns and chronicity. In this study, we examined the effects of CHD on blood monocytes and alveolar macrophages in control and CHD female macaques after 12 months of daily ethanol consumption. While monocytes from CHD female macaques generated a hyper-inflammatory response to ex vivo LPS stimulation, their response to *E*.*Coli* was dampened. In depth scRNA-Seq analysis of purified monocytes revealed significant shifts in classical monocyte subsets with accumulation of cells expressing markers of hypoxia (*HIF1A*) and inflammation (NFkB signaling pathway) in CHD macaques. The increased presence of monocyte subsets poised to generate a hyperinflammatory response was confirmed by the epigenetic analysis which revealed higher accessibility of promoter regions that regulate genes involved in cytokine signaling pathways. Finally, alveolar macrophages (AM) from the same animals produced higher levels of inflammatory mediators in response to LPS stimulation, but reduced ability to phagocytose bacteria. Collectively, data presented in this manuscript demonstrate that CHD primes monocytes and tissue-resident macrophages towards a more hyper-inflammatory immune response with compromised functional abilities, which could be used in diagnostic purposes or preventative measures for patients with alcohol use disorders.

## INTRODUCTION

Alcohol consumption is widespread in the United States with 85% of individuals ages 18 and older engaging in this behavior. While the overwhelming majority of these individuals are considered moderate drinkers, 7% are classified as heavy alcohol users (National Survey on Drug Use and Health, 2015). Chronic heavy alcohol consumption or chronic heavy drinking (CHD) is associated with multiple adverse health effects including increased incidence of cardiac disease (1, 2), certain types of cancer (3-6), liver cirrhosis (7), and sepsis (8), making it the third leading preventable cause of death in the United States (9). CHD is also associated with higher susceptibility to bacterial and viral infections including pneumonia and tuberculosis (10, 11), hepatitis C virus, and HIV (12, 13). Moreover, CHD compromises tissue repair, resulting in reduced post-operative healing and poor trauma recovery outcomes (14, 15). These observations strongly suggest that CHD dysregulates immunity and host defense.

While CHD can modulate the function of many immune cell populations, data from several laboratories indicate that the most dramatic and consistent changes are evident in the innate immune branch, specifically in myeloid cells (monocytes, macrophages, dendritic cells, and neutrophils) (16-18). Monocytes are relatively short-lived phagocytic cells that circulate in the blood, are constantly repopulated from bone marrow progenitors, and can quickly respond to infection or inflammation by extravasation into tissue and differentiation into tissue-resident macrophages (19). A tightly regulated inflammatory response by monocytes is required for effective infection clearance and tissue repair (20). Alcohol consumption has been shown to disrupt monocyte and macrophage responses in a dose and time dependent manner (16).

Specifically, acute alcohol treatment of purified primary human monocytes, rodent monocyte-derived macrophages, or human monocytic cell lines results in decreased production of inflammatory mediators including TNFα (16) following stimulation with lipopolysaccharide (LPS), a TLR4 agonist (21, 22). In rodents, acute *in vivo* (23) exposure to ethanol increases production of the anti-inflammatory cytokine IL-10 through activation of STAT3. Similarly, in healthy human subjects, an acute binge of alcohol increased IL-10 and decreased IL-1β production (24). In contrast, prolonged exposure to alcohol results in increased production of pro-inflammatory mediators, notably TNFα in response to LPS or PMA stimulation (22, 25) potentially due to enhanced activation of NFκB and ERK kinases (26). In line with these observations, monocytes as well as tissue resident macrophages, including Kupffer cells (27), microglia (28), alveolar macrophages (29), and splenic macrophages (17, 30) taken from patients with alcoholic liver disease (ALD) produce higher levels of TNF-α at resting state as well as in response to LPS (31). The enhanced inflammatory response by tissue-resident myeloid cells in the context of CHD is linked to organ damage, most notably in the liver (32) but also the intestine (33), brain (34, 35), and lungs (36).

Despite the studies described above, our understanding of the mechanisms by which chronic alcohol consumption reprograms circulating monocytes and tissue-resident macrophages remains incomplete. Some studies have suggested that CHD-induced gut “leakiness” and translocation of bacterial products including LPS across the gut barrier into the circulation could lead to chronic activation and subsequently organ damage (33). Whether monocytes in circulation are activated prior to organ damage by circulating bacterial products, ethanol, ethanol metabolites, or a combination of these factors remains unanswered.

Data from clinical studies are confounded by self-reported alcohol intake, the use of nicotine, recreational or illicit drugs, nutritional deficiencies, and presence of organ damage (37). Addressing these questions requires access to a reliable animal model that recapitulates critical aspects of human CHD. Therefore, to better model human CHD and relate immune response of peripheral monocytes and resident macrophages to quantified alcohol intakes in the absence of confounders listed above, we leveraged a rhesus macaque model of chronic voluntary ethanol self-administration (38, 39). Using this model, we reported that CHD disrupts the resting transcriptome and results in heightened inflammatory responses by peripheral blood mononuclear cells (PBMC) from male and female macaques (18, 40). Additionally, splenic macrophages from CHD monkeys generate a hyper-inflammatory response following LPS stimulation that is accompanied by increased chromatin accessibility at promoters and intergenic regions that regulate genes important for inflammatory responses (41).

In this study, we examine how CHD disrupts the inflammatory response of blood monocytes and alveolar macrophages. We show that CHD is associated with increased numbers of circulating monocytes that exhibit a heightened transcriptional and immune mediator response to LPS stimulation, but lower response to bacterial pathogens. To determine the molecular mechanisms of this dysregulated response, we profiled the transcriptome of the monocytes by single cell RNA-Seq and their epigenetic landscape by ATAC-Seq. Finally, we show that the heightened inflammatory response coupled by diminished anti-microbial responses extend to alveolar macrophages. Our results indicate that CHD significantly alters the epigenetic and transcriptional profiles of circulating monocytes, priming them towards a hyper-responsive state and altering their effector function in tissue.

## RESULTS

### Chronic heavy drinking-induced enhanced innate immune response in PBMC is independent of sex

Recent bulk RNA seq analysis of resting PBMC obtained from female rhesus macaques after 12 months of chronic heavy drinking (CHD) indicated that most of the differential gene expression originated from innate immune cells (monocytes and dendritic cells (DCs)) (14). Moreover, studies using PBMC from male macaques showed dysregulated response to ex vivo LPS stimulation with CHD after 12 months of chronic ethanol consumption (40). To assess if CHD also led to changes in inflammatory responses by circulating innate immune cells in female macaques, PBMC obtained from CHD (n=6) and control (n=3) females were stimulated ex vivo with LPS for 16 hours followed by measurement of immune mediator production by Luminex assay and transcriptional changes by bulk RNA Seq (**Supp. Figure 1A**). LPS stimulation resulted in robust inflammatory response (TNFα, IL-6, IL-18, IL-4, IL-8, GMCSF and S100B) by PBMC from both CHD and control animals (**Figure 1A**). However, PBMC from CHD animals produced higher levels of additional inflammatory mediators notably IL-1β, IL-12, IFNγ, CCL3, and CCL4 (**Figure 1A**). Moreover, production of inflammatory cytokines IL-6, IL-23 and to lesser extent soluble PD-L1 were higher in CHD PBMC following LPS stimulation compared to controls (**Figure 1A**). In addition, levels of TNFα, CCL4, IL-6, IL-15, IL-23, and sPD-L1 were significantly positively correlated with ethanol dose (**Figure 1B and Supp. Figure 1B**).

**Figure 1:**
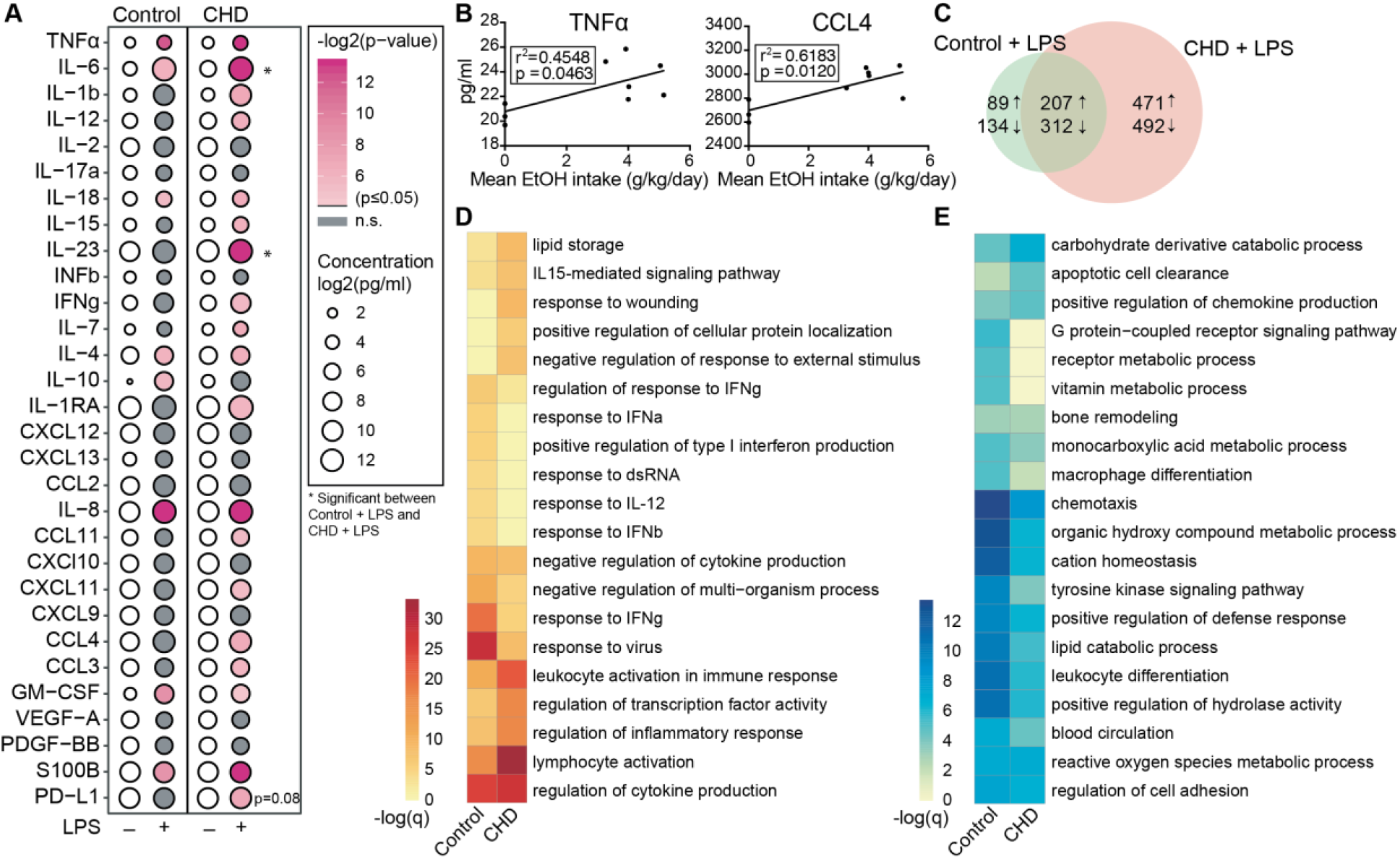
CHD induced enhanced innate immune response in the periphery. PBMC from control (n=3) and chronic heavy drinking (CHD) (n=6) animals were stimulated with LPS for 16 hours. A) Bubble plot representing immune factor production (pg/ml) in the presence or absence of LPS stimulation of PBMC from control and CHD animals. The size of each circle represents the log_2_ mean concentration of the indicated secreted factor and the color denotes the -log_2_ transformed p value with the darkest pink representing the most significant value. The p-values were calculated between the unstimulated and stimulated conditions for each group by One-way ANOVA and a p-value cut-off of 0.05 was set. White circles indicate non-significant p-value. * indicates significance between control and CHD for each stimulation condition. B) Scatter plots showing Spearman correlation between average EtOH dose (grams EtOH/kg body weight/day) and concentration (pg/ml) of the secreted factors TNFα and CCL4. C) Venn diagram comparing LPS-induced DEG in controls and CHD PBMC. D, E) Heatmaps of significant GO terms to which upregulated (D) and downregulated (E) DEG identified following LPS stimulation of PBMC from controls and CHD animals enriched using Metascape. The scales of the heatmaps are -log(q-values) associated with the enrichment to selected pathways.

We next assessed transcriptional differences in response to LPS between the two groups. Differential analysis indicated a heightened response to LPS by CHD PBMC relative to healthy control PBMC (**Figure 1C**). While signatures of immune activation were observed in both groups following LPS exposure, DEGs upregulated in the controls only enriched to pathways involved in antiviral immunity (“response to IFNγ” and “response to virus”), whereas DEGs upregulated only in PBMC from CHD animals enriched to leukocyte activation and inflammation pathways (**Figure 1D and Supp. Figure 1C**). Enrichment of the downregulated genes shows significant enrichment to GO terms associated with chemotaxis, leukocyte activation and metabolism in the PBMC from control animals (**Figure 1E and Supp. Figure 1D**).

### Chronic heavy drinking impacts monocyte frequencies and functional responses to LPS stimulation

To measure changes in frequencies and phenotypes of monocyte subsets with CHD, we profiled PBMC by flow cytometry. Interestingly, the proportions of monocytes within PBMC from CHD macaques were significantly elevated, however the relative distribution of the monocyte sub-populations were comparable between the groups (**Figure 2A and Supp. Figure 2A, B**). We profiled a number of TLRs, chemokine, and activation receptors on the monocytes by flow cytometry, but found no significant differences (**Supp. Figure 2C**). To assess whether the heightened inflammatory response detected in PBMC was due to increased numbers of monocytes or cell-intrinsic changes caused by CHD, monocytes were purified from each group and subjected to bulk RNA-Seq at resting state and after LPS stimulation (**Supp. Figure 1A**). A modest number of DEG were detected at resting state (58 upregulated and 112 downregulated) between CHD and control monocytes (**Figure 2B**). The DEG upregulated with CHD enriched to pathways associated with inflammatory response such as “pattern recognition receptor activity” (e.g. *NLRP3*) and “cytokine production” (e.g. *FCN1)* (**Figure 2C, and Supp. Figure 2D)**. DEG downregulated with CHD mapped to gene ontology (GO) terms “activation of immune response” (e.g. *IFNG*) “lymphocyte proliferation” (e.g. *HSPD1*) and “chemotaxis” (e.g. *CCR7*) (**Figure 2C**). One potential mechanism for this enhanced monocyte transcriptional activation at resting state could be increased microbial products in circulation due to altered barriers (31). Therefore, we measured circulating levels of bacterial endotoxin (LAL) and IgM bound endotoxin. We found slightly increased levels of circulating LAL with CHD, but no changes in IgM-bound endotoxin levels (**Supp. Figure 2E, F**).

**Figure 2:**
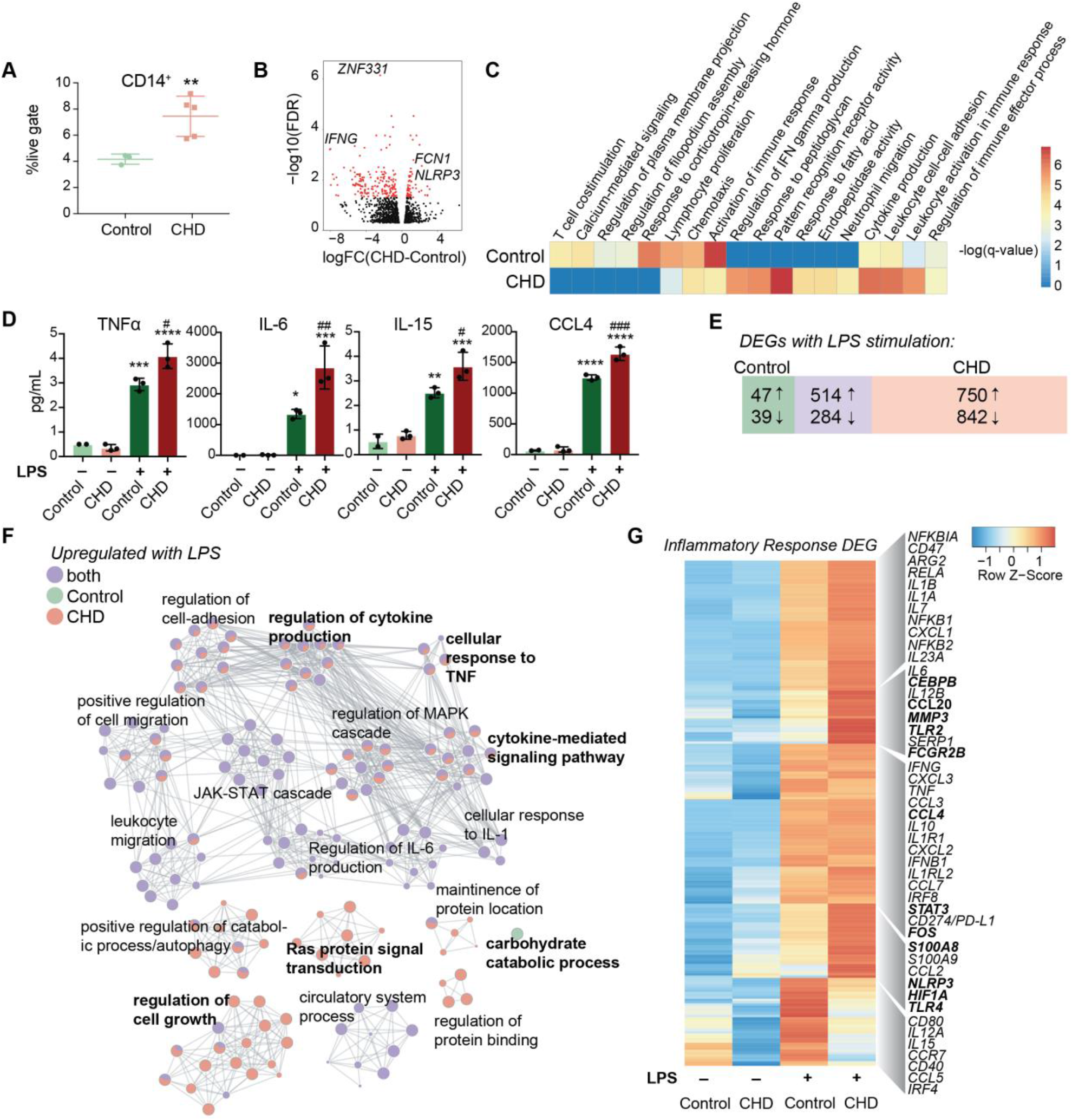
Impact of CHD on the monocyte functional and transcriptomic response to LPS. A) Abundance of live CD14+ cells in PBMC from control (n=3) and CHD (n=5) animals measured by flow cytometry. Significance calculated by t-test with Welch’s correction. B) Volcano plot representing up- and downregulated differentially expressed genes (DEG) with CHD in resting monocytes. Red = significant with an FDR ≤ 0.05 and fold change ≥ 2. C) Heatmap of significant GO terms (identified using Metascape) to which DEGs downregulated and upregulated with CHD in monocytes enriched. The scales of the heatmaps are -log(q-values) associated with the enrichment to selected pathways. D) Purified monocytes were stimulated with LPS for 6 hours and supernatants were analyzed by Luminex assay. Bar plots showing concentration (pg/ml) of selected immune mediators (TNFα, IL-6, IL-15, CCL4) (*=p<0.05, **=p<0.01, ***=p<0.001, ****=p<0.0001 between LPS and unstimulated condition, # is significance between CHD and control for each stimulation condition calculated by One-way ANOVA). E) Venn diagram showing overlap between DEG identified in control and CHD monocytes after LPS stimulation. F) Network visualization of functional enrichment analysis of DEG upregulated with LPS in monocytes from controls only, CHD only, and both using Metascape. The size of each node represents the number of DEGs associated with a given gene ontology (GO) term and the pie chart filling represents relative proportion of DEGs from each group that enriched to that GO term. Similar GO terms are clustered together and are titled with the name of the most statistically significant GO term within that cluster. The gray lines denote shared interactions between GO terms. Density and number of gray lines indicates the strength of connections between closely related GO terms. G) Heatmap of average normalized expression of genes associated with inflammatory response pathways.

Secreted levels of cytokines, chemokines, and growth factors were measured after 6-hour incubation in the presence or absence of LPS by Luminex assay. No significant differences in the concentration of immune mediators were noted in the non-stimulated conditions. As observed for PBMC, significantly enhanced production of pro-inflammatory cytokines TNFα, IL-6, and IL-15, chemokines CCL4, and CXCL11, and to a lesser extent IL-4 and IL-7 by CHD monocytes were noted (**Figure 2D and Supp. Figure 3A)**. The LPS response was also assessed at the transcriptional level using RNA-Seq. Principal component analysis (PCA) showed that stimulation accounts for the majority of transcriptional changes (PC1, 75% of the differences) (**Supp. Figure 3B**) with 357 and 262 DEG between CHD and controls in the NS, and LPS conditions, respectively (**Supp. Figure 3C, D**). The DEG from these comparisons were similar to those detected in resting cells, with genes associated with myeloid inflammatory pathways are upregulated (*TNFSF21, MMP2, TLR2, MMP1*) while genes associated with adaptive immune activation are downregulated (*IL21R, CD40, MAMU-DOA, CCR6*) in CHD compared to control monocytes (**Supp. Figure 3C, D**).

We then identified DEG in the LPS relative to NS condition for each group. A greater number of DEG was detected in the CHD group, with a large overlap between the two groups, and few DEG detected solely in the control group (**Figure 2E**). As expected, DEG common between the two groups enriched to GO terms associated with monocyte activation including “regulation cytokine production”, “leukocyte migration”, and “JAK-STAT cascade” (**Figure 2F**). The DEG detected only in control animals mapped to “Carbohydrate catabolic process”. The DEG unique to monocytes from CHD animals mapped to cytokine production and signaling pathways (**Figure 2F**). The expression of inflammatory genes was broadly upregulated in the CHD monocytes following LPS stimulation including *CEBPB, CCL20, CCL4, STAT3, FOS, S100A8, HIF1A*, and *TLR4* (**Figure 2G**). Additional predictive analysis revealed that LPS-responsive DEGs detected in the CHD monocytes are regulated by canonical transcription factors NFKB2, RELB, and JUNB with over 250 genes mapping to each (**Supp. Figure 3E**).

### Chronic heavy drinking alters the monocyte transcriptome and cell subset distribution at the single cell level

To gain a deeper understanding of the heightened activation state of monocytes with CHD, we performed single cell RNA sequencing (scRNA-Seq) on sorted CD14+HLA-DR+ monocytes from CHD and control female macaques (n=3 pooled samples/group). UMAP clustering of all 9,360 monocytes revealed 9 distinct monocyte subsets (MS) (**Figure 3A**). These clusters could be categorized into the three major monocyte subtypes typically identified by flow cytometry: non-classical, intermediate, and classical based on expression of *CD14, MAMU-DRA*, and *FCGR3*(CD16) (**Supp. Figure 4A**). As we identified by flow cytometry, the frequency of these three major subsets was comparable between controls and CHD animals by scRNA-Seq analysis (**Supp. Figure 4B**). The non-classical monocytes formed one cluster (MS6) exhibiting high expression of *CX3CR1, MS4A7*, and *FCGR3* and lower expression of *CD14* and MHC II molecule *MAMU-DRA* (**Figure 3A, B, and Supp. Figure 4C**). The intermediate monocytes also fell into one cluster (MS4), expressing lower levels of non-classical (*MS4A7, FCGR3A*) and classical (*CD14, S100A8*) markers with higher expression of *MAMU-DRA* (**Figure 3A, B, and Supp. Figure 4C**). Finally, 7 clusters of classical monocytes were identified (MS1, MS2, MS3, MS5, MS7, MS8, and MS9), based on expression of *CST3, GOS2, S100A8, S100A9, HIF1A, SOD2, EGR1*, and *HERC5* (**Figure 3A, B, and Supp. Figure 4C**). These data reveal considerable, previously unappreciated transcriptional heterogeneity within the classical monocytes. Subsets MS5, MS7 were primarily detected in monocytes from CHD animals while MS8 was primarily detected in monocytes from control animals (**Figure 3 and Supp. Figure 4D**). Module scoring revealed that MS5 had the lowest expression of genes that play a role in antigen presentation as well as hypoxia, while MS5 and MS7 had higher expression of genes within the TLR-and IL-6-signaling pathways (**Supp. Figure 4E**).

**Figure 3:**
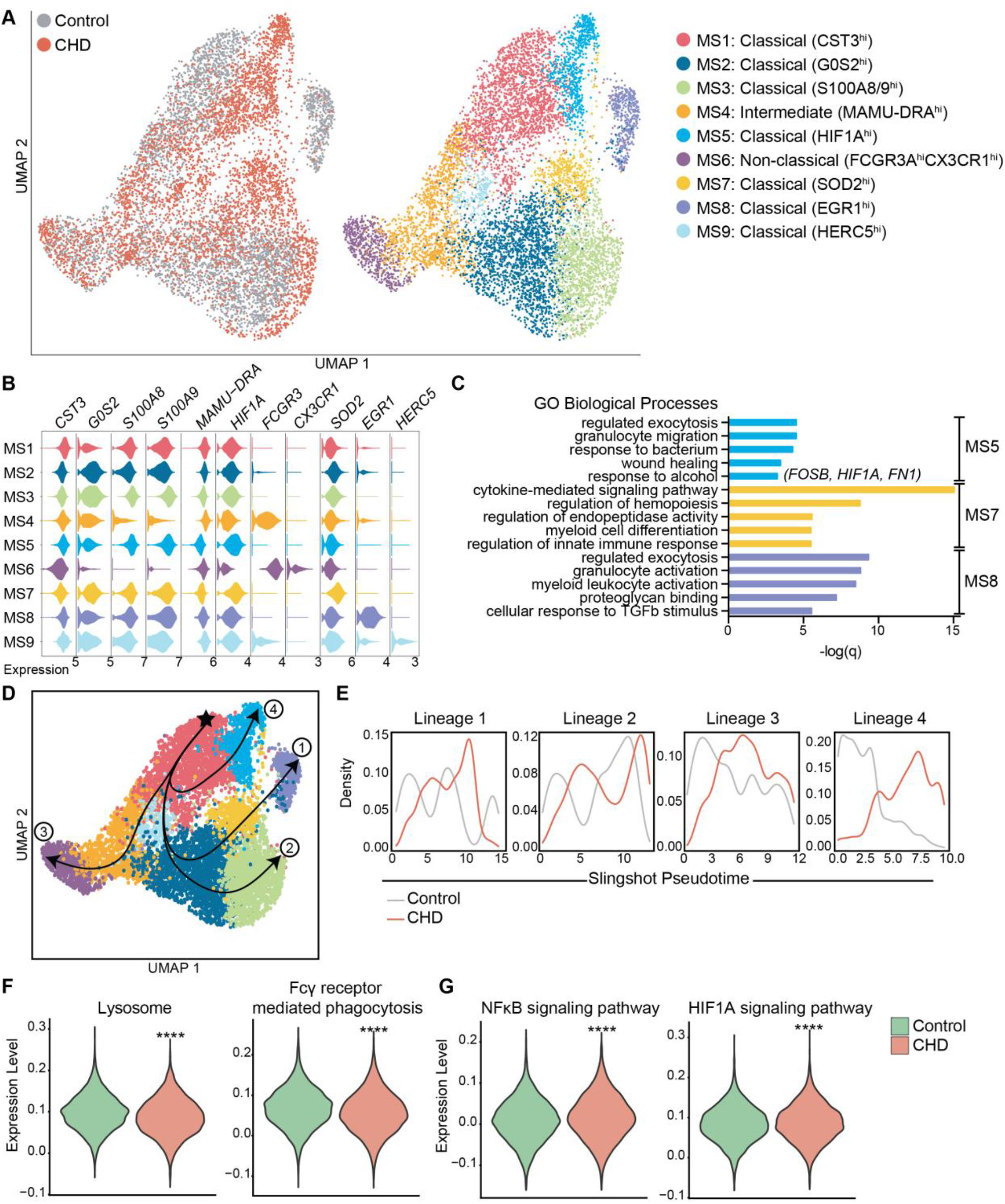
Single cell RNA-Seq analysis of monocytes with CHD. Monocytes (n=3 control/ 3 CHD) were purified from total PBMC and subjected to 10X scRNA-Seq analysis. A) Visualization of 9,360 monocytes by uniform manifold approximation and projection (UMAP) colored by group (control and CHD) as well as by identified clusters. B) Stacked violin plots representing expression of key genes used for cluster identification, grouped by monocyte subset cluster. C) GO Biological Process enrichment from Metascape of genes highly expressed in MS5, MS7, and MS8, defined by q-value. D) Trajectory analysis of monocytes determined using Slingshot. E) Cell density plots for Control and CHD groups across each of four trajectory lineages determined by Slingshot. F,G) Violin plots comparing (F) Lysosome and Fcγ receptor-mediated phagocytosis and (G) NFkB signaling and HIF1A signaling pathway module scores in all monocytes of both groups. Statistical analysis of module scores was performed using a t-test with Welch’s correction where *=p<0.05, **=p<0.01, ***=p<0.001, ****=p<0.0001.

To identify the biological implications of these subsets, we extracted the genes that define each cluster and performed functional enrichment (**Figure 3C**). Although genes that define MS5 (classical – CHD) and MS8 (classical – control) subsets enriched to similar GO terms (“Regulated exocytosis”, “granulocyte activation/migration”), only genes highly expressed in MS5 (CHD animals) enriched to “Response to alcohol” and “Wound healing”, notably *FOSB, HIF1A*, and *FN1*. Interestingly, genes that define MS7 subset (classical – CHD) enriched to “Cytokine-mediated signaling pathway” including *CSF3R, IRF1, IRF7, NFKBIA, STAT1, STAT3*, and *VEGFA*. Classical subsets MS1 and MS2 were slightly more abundant in monocytes from control animals and expressed high levels of *CST3* and *GOS2* as well as *S100A8* and *S100A9*, respectively.

To better understand the relationships between the classical monocyte clusters and their differentiation/activation states, we performed a trajectory analysis. The monocyte clusters were ordered by pseudotime, starting with the most abundant MS1 cluster which resulted in four unique trajectory lineages (**Figure 3D**). Lineages 1, 2 and 4 indicate transitions culminating in MS8, MS3, and MS5, respectively. Lineage 3 is the classical to non-classical differentiation trajectory. Density plots revealed enrichment of control monocytes at the start of each lineage (less differentiated) and enrichment of the CHD monocytes at the end of the lineages (more differentiated), most notably in Lineage 4 (**Figure 3E**).

We next assessed differential gene expression with CHD in the 3 major subsets (**Supp. Figure 4F**). This analysis showed that IFN-inducible genes *WARS* and *IDO* were upregulated in the classical CHD monocytes, whereas MHC-II gene *MAMU-DRB1* was downregulated (**Supp. Figure 4F**). In the intermediate monocytes, CHD induced upregulation of inflammatory signaling genes MAP3K and S100P but led to decreased expression of FC-receptor gene *FCGR3* (**Supp. Figure 4F**). The 33 DEG downregulated in the non-classical monocytes from the CHD group relative to controls enriched to antigen processing and IFNγ signaling pathways (**Supp. Figure 4G**). Finally, broadly in all monocytes we identify a significant reduction in lysosome and Fcγ-receptor mediated phagocytosis modules in the CHD monocytes (**Figure 3F**). Alternatively, the CHD monocytes show increased expression of gene modules for NFkB signaling and HIF1A signaling (**Figure 3G**).

### Chronic heavy drinking impacts the epigenome of circulating monocytes

To uncover epigenetic basis for the altered transcriptional profile in resting monocytes and altered functional responses with CHD, we profiled the chromatin accessibility in purified resting monocytes using the assay for transposase-accessible chromatin sequencing (ATAC-Seq) (42). We identified 11,717 differentially accessible regions (DAR) that were open in the CHD monocytes compared to 9,173 DAR that were accessible in the controls (**Figure 4A**). More than 25% of the accessible regions in the CHD monocytes mapped to promoter regions and were closer to the transcription start site compared to accessible regions in monocytes from control animals (**Figure 4A and Supp Figure 5A**). Genes regulated by promoter regions open in the CHD monocytes enriched to GO terms associated with cytokine production and myeloid cell activation (**Figure 4B**). Additional analysis using GREAT showed that cis-regulatory elements in the distal intergenic regions open in CHD monocytes enriched to processes involved with apoptotic signaling, MAPK signaling cascade, and myeloid cell differentiation (**Supp Figure 5B**). Motif enrichment analysis of open chromatin regions in the CHD and control monocytes demonstrated increased putative binding sites for transcription factors important for monocyte differentiation and activation SP1, FRA1, FOS, JUNB, BATF, and PU.1 with CHD (43-47)(**Figure 4C**). To link the resting epigenome of the CHD monocytes to the enhanced transcriptional response after LPS stimulation, we compared the genes associated with the open promoter regions (<1kb) and the DEG identified after LPS stimulation and identified 281 common genes (**Figure 4D**). Functional enrichment of these genes showed significant mapping to Biological Process “Cytokine-mediated signaling” including transcription factors *FOS, HIF1A, STAT3* and *JUNB* (**Figure 4E**). These genes as well as inflammatory *CCL20, TLR2, CD81* and *TNFSF14* had both open promoters in the resting CHD monocytes as well as upregulated gene expression in the LPS-stimulated CHD monocytes, linking open chromatin under resting conditions with CHD to transcriptional LPS responses (**Figure 4F**).

**Figure 4:**
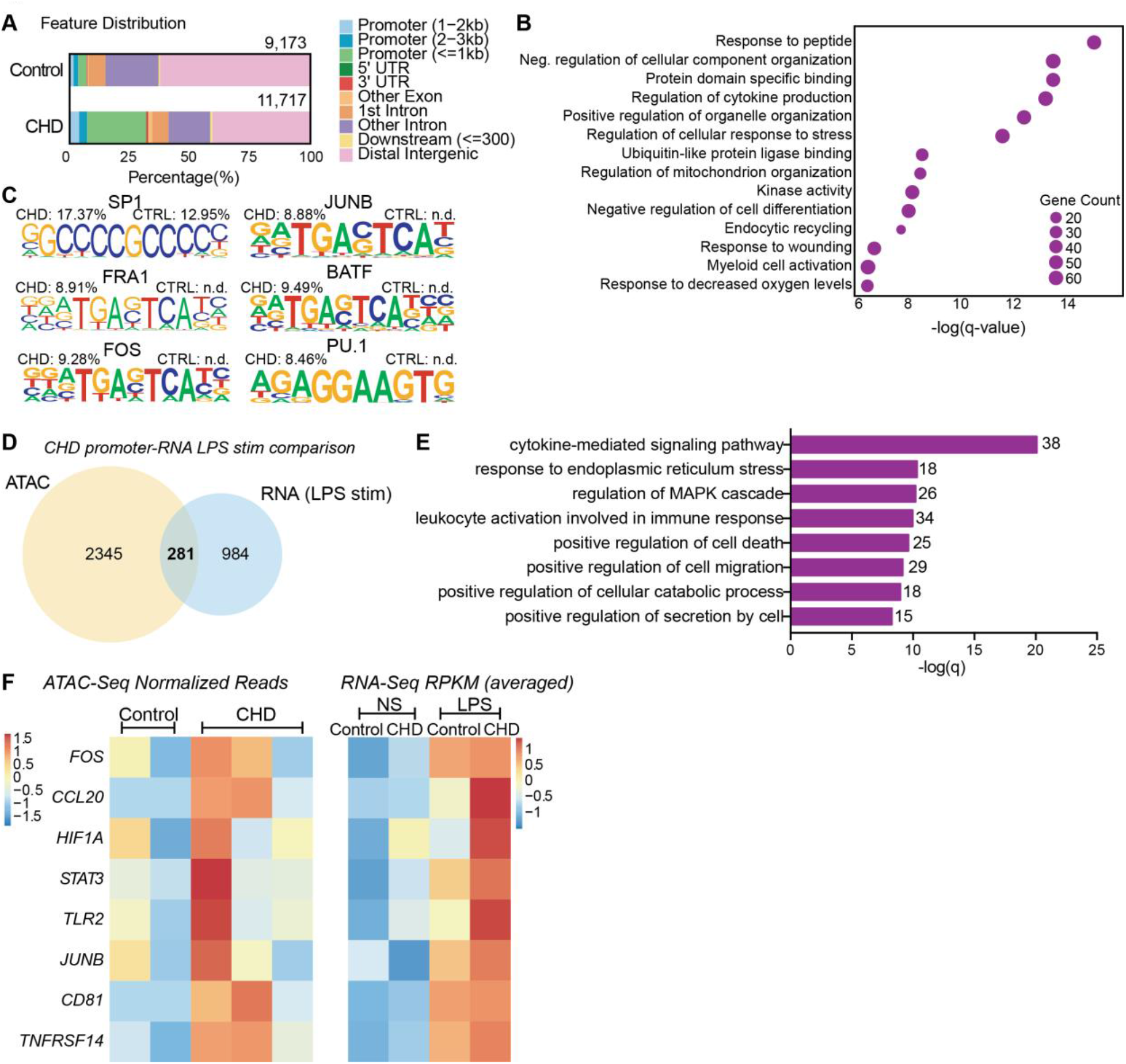
CHD primes the monocyte epigenome for heightened response. Monocytes (n= 2 Control/ 3 CHD) were purified from PBMC and subjected to ATAC-Seq analysis. A) Bar plot showing genomic feature distribution of the open chromatin regions (fold-change ≥ 2) in control and CHD monocytes at resting state. B) GO Biological process enrichment from Metascape of genes regulated by the open promoter regions (≤ 1kb, fold-change ≥ 3) in CHD monocytes. The X-axis represents -log(q-value) and the size of the dot represents the number of genes within that term. C) Homer motif enrichment of the open chromatin regions. All listed motifs have significantly enriched binding sites in the CHD monocytes where the percentage value listed is the percentage of target sequences with that motif. D) Venn diagram of genes regulated by the open promoter regions (≤ 1kb, fold-change ≥ 3) and DEG detected following LPS stimulation in the CHD monocytes. E) GO Biological process enrichment terms from Metascape of the 281 overlapping genes from (D). F) Heatmaps of the normalized expression of open promoter region counts and RPKM from bulk RNA-Seq analysis for selected common genes.

### CHD impairs monocyte response to E. coli

We next asked whether CHD would affect the monocyte response to pathogens. Monocytes from male and female macaques were co-cultured with heat-killed *E*.*coli* for 16 hours and immune mediator production was measured using Luminex (**Supp. Figure 1A**). In contrast to the response to purified LPS, monocytes from CHD animals generated a dampened response to *E*.*coli* characterized by reduced levels of IL-1b, IL-6, MIP-1B, and IL-15 and to a lesser extent I-TAC and IL-5 compared to their control counterparts (**Figure 5 A, B**).

**Figure 5:**
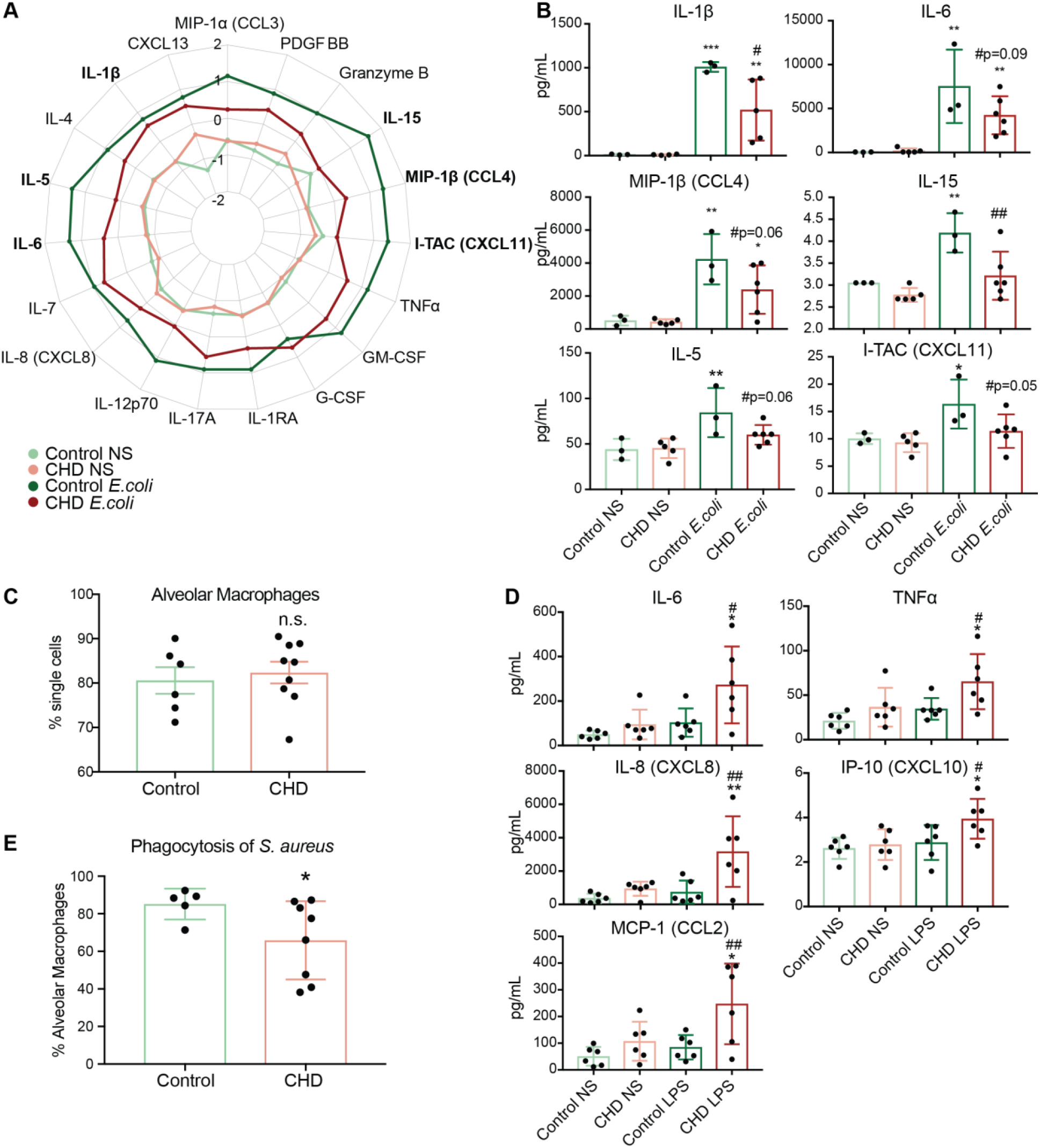
Alveolar macrophage function and response to LPS is altered with CHD. Purified monocytes (n= 3 Control/ 5 CHD) were stimulated with heat-killed *E*.*coli* bacteria for 16 hours. A) Spider plot representing average Z-scores for each group across the indicated analytes measured by Luminex. B) Bar plots showing concentration (pg/ml) of selected immune mediators (TNFα, IL-6, MIP-1b, IL-15, IL-5, I-TAC).C) Flow cytometry analysis of live CD206+ cells from the BAL of macaques. D) FACS sorted AM (n=6/group) were stimulated with LPS for 16 hours and production of IL-6, TNFα, IL-8, IP-10, and MCP-1 were quantified by Luminex assay. E) Bar plots showing percentage of AM phagocytosing fluorescent *S. aureus* bacteria. Statistics for 2-way comparisons carried out using t-test with Welch’s correction, 4-way by One-way ANOVA. *=p<0.05, **=p<0.01, ***=p<0.001, ****=p<0.0001 between LPS and unstimulated condition. If indicated, # is significance between CHD and control for each stimulation condition.

### Tissue-resident macrophage function is disrupted with CHD

To understand functional consequences of CHD in relevant tissue-resident macrophages, we assessed alveolar macrophages (AM) isolated from bronchoalveolar lavage (BAL) samples taken from male and female macaques. Unlike our results from blood (**Figure 2A**) and the spleen (41) myeloid cells, no differences in relative frequency of AM were noted in the BAL with CHD (**Figure 5C**). Purified AM from control and CHD animals were stimulated with LPS (16 hours) and production of immune mediators was measured by Luminex (**Supp. Figure 1A**). In contrast to AM from control animals, which did not mount a significant response to the LPS, AM from CHD animals secreted large amounts of inflammatory cytokines IL-6 and TNFα as well as chemokines IL-8, IP-10 and MCP-1 (**Figure 5D**). Finally, we assessed the impact of CHD on phagocytic ability of AM (**Supp. Figure 1A**). AM from CHD macaques had a reduced ability to phagocytose *S. aureus* bacteria (**Figure 5E**).

## DISCUSSION

It is well-appreciated that chronic alcohol drinking exerts a profound impact on peripheral and tissue-resident innate immune cells. However, the limitations provided by the complexity of studying alcohol consumption in humans has left major gaps in understanding mechanisms that underlie the immune response under conditions of heavy alcohol drinking. In this study, we utilized a macaque model of voluntary ethanol self-administration to profile peripheral monocytes and tissue-resident macrophages in animals after 12 months of daily alcohol drinking. We first demonstrate that circulating monocytes from CHD animals generate a hyper-inflammatory response to ex vivo LPS stimulation at the transcript and protein level. A comprehensive profiling of circulating monocytes by scRNA-Seq and ATAC-Seq revealed alterations in monocyte differentiation state as well as the epigenetic landscape with CHD. In contrast to what we observed following ex vivo stimulation with LPS, monocytes from CHD animals generated a dampened production of immune mediators following ex vivo *E*.*coli* stimulation. This dysregulated phenotype extended to alveolar macrophages (AM) obtained from bronchia alveolar lavage (BAL) samples which generated a heighted response to LPS, but reduced bacterial phagocytosis.

The enhanced inflammatory responses in response to ex vivo stimulation with LPS by PBMC from CHD female macaques are in line with our previous data for PBMC from CHD male macaques (40), indicating that this consequence of CHD is sex independent. Importantly, our analysis indicated that this hyper-inflammatory response is mediated largely by intrinsic changes within monocytes. Indeed, CHD induced significant transcriptional differences consistent with activation of inflammatory pathways in monocytes at resting state. Moreover, LPS stimulation resulted in a drastic increase in numbers of DEG from the CHD monocytes with increased expression of genes associated with inflammatory response. In contrast, genes involved in carbohydrate catabolic processes were not upregulated in monocytes from CHD animals suggesting metabolic rewiring associated with a heightened inflammatory state (48). The increased inflammatory mediator production in response to ex vivo LPS stimulation are in line with previous *in vitro* studies that have reported hyper-activation of myeloid cells with prolonged alcohol exposure. This also fits with hyper-inflammatory phenotypes associated with patients with alcohol use disorders, especially those with alcoholic liver disease (16, 26, 49).

To elucidate the molecular basis for this dysregulated inflammatory response, we profiled monocytes from CHD and control animals by scRNA-Seq. This analysis revealed tremendous diversity amongst non-classical monocytes that has not been previously appreciated. We identified two unique classical clusters within monocytes from CHD animals expressing high levels of hypoxia factor *HIF1A* (MS5), and antioxidant defense molecule *SOD2* (MS7). Alcohol and the products of its metabolism can induce oxidative stress which can alter cellular transcriptional profiles, potentially explaining the profile of the MS5 cluster (50). CHD animals also had a higher number of cells within the anti-viral cluster MS9 that expressed high levels of interferon stimulated (ISG) genes such as *HERC5*. This observation is in line with our earlier study that reported higher levels of ISG within PBMC from these same animals (18). Functional module scoring of all of the monocytes revealed downregulation of genes involved in lysosome function and Fcγ receptor mediated phagocytosis, but upregulation of NFkB signaling, HIF1A signaling and Fatty acid metabolism. Moreover, trajectory within the monocyte subsets revealed that CHD accelerates differentiation/activation of classical monocyte subsets making them potentially poised towards a hyper-inflammatory response.

Although the non-classical and intermediate monocytes fell into single clusters with equal numbers of cells from controls and CHD animals, differential gene expression analysis revealed significant downregulation of genes associated with MHC class II, antigen processing and presentation, and IFNγ signaling pathways in the CHD monocytes. This observation provides a potential explanation for the reduced response to vaccination observed in CHD animals (51). Indeed, monocytes from CHD animals generated a dampened cytokine and chemokine response to heat killed *E*.*coli*. These observations are in line with reported increased susceptibility of patients with alcohol use disorders to bacterial pathogens (10, 11). Collectively, these data suggest a rewiring of monocytes with CHD towards inflammatory responses and away from anti-microbial functions in additional to alterations in signaling and metabolic processes.

Ethanol metabolites notably acetaldehyde and acetate in addition to minor byproducts such as reactive oxygen species (ROS) and lipid peroxidation products can modulate gene expression by binding transcription factors and/or modifying chromatin accessibility (50). Therefore, we profiled chromatin accessibility of monocytes from control and CHD animals by ATAC-Seq. We discovered that promoter regions that regulate genes important for cytokine production and myeloid cell activation were more accessible in monocytes from CHD animals. These open regions contained putative binding sites for transcription factors important for monocyte activation and differentiation. Integrative analyses showed correlation between chromatin accessibility and expression levels of key inflammatory genes in CHD monocytes, including *FOS, HIF1A*, and *JUNB*.

To assess the implications of our observations in CHD monocytes in a functionally relevant tissue, we profiled alveolar macrophages (AM) in the same animals. While it is believed that there is a population of yolk-sac derived lung macrophages with self-renewal potential, there are also populations of lung macrophages that arise from infiltrating monocytes (52). Similar to monocytes, CHD AM produced significantly more inflammatory TNFα, IL-6 and other chemokines following ex vivo LPS stimulation. Additionally, AM from CHD animals were compromised in their ability to phagocytose *S. aureus* compared to controls as has been described in rodent models of acute alcohol exposure in rodents (53-55). These data indicate that the mis-wiring we observed in circulating monocytes extends to tissue residence macrophages.

One proposed mechanism of this hyper-inflammatory state in monocytes (32, 34-36) and macrophages with CHD is the translocation of bacterial products into circulation through impaired gut barrier caused by ethanol consumption (33). In a previous study, we report increased levels of IgM-bound endotoxin in CHD in male macaques (56). Here, we detected modestly elevated levels of LAL in circulation but no changes in IgM bound endotoxin. A small increase in these circulating bacterial products could significantly impact the activation state of monocytes; however, it is difficult to tease these effects out from circulating ethanol and its metabolic products (acetaldehyde, acetate). Long term exposure to activating agents, like LPS, is believed to lead to tolerance and a decreased response to secondary stimulation in monocytes and macrophages defined by specific changes in chromatin structure (57-60). Interestingly, we see enhanced inflammatory responses of monocytes and macrophages from CHD animals, akin to innate immune training (61, 62). Indeed, ethanol and its metabolites have been found to directly act on histone modifications, creating changes in accessible chromatin (50, 63, 64). This could be occurring in circulating monocytes, tissue macrophages, or the progenitors of these cells in the bone marrow; perhaps on all three levels, altering the epigenetic structure and therefore functional capacities of these cells (65).

In summary, this study provides a novel, in-depth, and integrative analysis of the effects of long-term *in vivo* alcohol drinking on myeloid cells in non-human primates. To the best of our knowledge, this is the first study to examine the effects of CHD on blood monocytes and tissue-resident macrophages from the same subjects. Additionally, the comprehensive analysis of monocyte populations alone by scRNA-Seq was critical for probing the heterogeneity of the classical monocyte compartment, which had not yet been appreciated in humans or macaques. Future studies will investigate the mechanisms behind the increased chromatin accessibility, including the role of histone modifications and transcription factor binding and whether epigenetic changes are apparent in the alveolar space as well.

## METHODS AND MATERIALS

### Animal studies and sample collection

These studies used samples from a non-human primate model of voluntary ethanol self-administration established through schedule-induced polydipsia (38, 39, 66). Briefly, in this model, rhesus macaques are introduced to a 4% w/v ethanol solution during a 90-day induction period followed by concurrent access to the 4% w/v solution and water for 22 hours/day for one year. During this time, the macaques adopt a stable drinking phenotype defined by the amount of ethanol consumed per day and the pattern of ethanol consumption (g/kg/day) (38). Blood samples were taken from the saphenous vein every 5-7 days at 7 hrs after the onset of the 22 hr/day access to ethanol and assayed by headspace gas chromatography for blood ethanol concentrations (BECs).

For these studies, blood and bronchoalveolar lavage (BAL) samples were collected from 9 female and 8 male rhesus macaques (average age 5.68 yrs), with 7 animals serving as controls and 10 classified as chronic heavy drinkers (CHD) based on 12 months of ethanol self-administration (tissue and drinking data obtained from the Monkey Alcohol Tissue Research Resource: www.matrr.com). These cohorts of animals (Cohorts 6 and 7a on matrr.com) were described in two previous studies of innate immune system response to alcohol (18, 40). Peripheral Blood Mononuclear Cells (PBMC) were isolated by centrifugation over histopaque (Sigma, St Louis, MO) as per manufacturer’s protocol and cryopreserved until they could be analyzed as a batch. BAL cells were obtained after 12 months of open access and centrifuged, pelleted and cryopreserved until they could be analyzed as a batch. The average daily ethanol intake for each animal is outlined in **Supp. Table 1**.

### LAL and IgM assays

Endotoxin-core antibodies in plasma samples were measured using an enzyme-immunoassay technique (ELISA) after 12 months of alcohol consumption using EndoCab IgM ELISA kit (Hycult Biotech, Catalog# HK504-IgM). Plasma samples were diluted 50x.

Circulating endotoxin was measured from plasma using a Limulus amebocyte lysate (LAL) assay (Hycult Biotech) following the manufacturers protocol.

### Flow cytometry analysis

1-2×10^6^ PBMC were stained with the following surface antibodies (2 panels) against: CD3 (BD Biosciences,SP34), CD20 (Biolegend, 2H7), HLA-DR (Biolegend, L243), CD14 (Biolegend, M5E2), CD16 (Biolegend, 3G8), TLR4 (Biolegend, HTA125), TLR2 (Biolegend, TLR2.1), CD40 (Biolegend, 5C3), CD163 (Biolegend, GHI/61), CD86 (Biolegend, IT2.2), CD80 (Biolegend, 2D10), CX3CR1 (Biolegend, 2A9-1), CCR7 (Biolegend, GO43H7), and CCR5 (Biolegend, J418F1) and live/dead Sytox Aadvanced (Invitrogen). Monocytes were defined as CD3-CD20-HLA-DR+CD14+. All samples were acquired with an Attune NxT Flow Cytometer (ThermoFisher Scientific, Waltham, MA) and analyzed using FlowJo software (Ashland, OR). Median Fluorescence Intensities (MFI) for all markers within the CD14+ monocyte gate were tested for significant differences using an unpaired t-test with Welch’s correction on Prism 7 (GraphPad, San Diego, CA).

### CD14 MACS bead Isolation and Purity

CD14+ monocytes were purified from freshly thawed PBMC using CD14 antibodies conjugated to magnetic microbeads per manufacturers recommendations (Milyenyi Biotec, San Diego, CA). Efficiency of the positive selection of monocytes was assessed by flow cytometry where purity (CD14+HLA-DR+) averaged 87% (SEM ± 1.6).

### Monocyte/Macrophage Stimulation Assays

1×10^6^ freshly thawed PBMC or 1×10^5^ purified CD14+ monocytes were cultured in RPMI supplemented with 10% FBS with or without 100 ng/mL LPS (TLR4 ligand, *E*.*coli* 055:B5; Invivogen, San Diego CA) for 16 or 6 hours, respectively, in 96-well tissue culture plates at 37C in a 5% CO_2_ environment. 3.5×10^4^ purified CD14+ monocytes were cultured in RPMI supplemented with 10% FBS with or without 6×10^5^ cfu/mL heat-killed *E*.*coli* (Escherichia coli (Migula) Castellani and Chalmers ATCC 11775) for 16 hours in 96-well tissue culture plates at 37C in a 5% CO_2_ environment. 6.5×10^4^ FACS sorted CD206+ cells from the BAL were cultured in RPMI supplemented with 10% FBS with or without 100 ng/mL LPS for 16 hours, in 96-well tissue culture plates at 37C in a 5% CO_2_ environment. Plates were spun down: supernatants were used to measure production of immune mediators and cell pellets were resuspended in Qiazol (Qiagen, Valencia CA) for RNA extraction. Both cells and supernatants were stored at −80C until they could be processed as a batch.

### Luminex Assay

Immune mediators in the supernatants from PBMC or purified monocytes stimulated with LPS were measured using a 30-plex panel measuring levels of cytokines (IFNγ, IL-1b, IL-2, IL4, IL-6, IL-7, IL-12, IL-15, IL-17, IL-18, IL-23, TNFα, IL-1RA, IFN-b, and IL-10), chemokines (MCP-1, MIP-1a, MIP-1b, Eotaxin, IL-8, MIG, I-TAC, BCA-1, IP-10, and SDF-1a), and other factors (PD-L1, PDGF-BB, S100B, GM-CSF, and VEGF-A) validated for NHP (R&D Systems, Minneapolis, MN, USA). Standard curves were generated using 5-parameter logistic regression using the xPONENT™ software provided with the MAGPIX instrument (Luminex, Austin TX).

Immune mediators in the supernatants from monocytes stimulated with *E*.*coli* or AM stimulated with LPS were measured using a more sensitive ProcartaPlex 31-plex panel measuring levels of cytokines (IFNα, IFNγ, IL-1β, IL-10, IL-12p70, IL-15, IL-17A, IL-1RA, IL-2, IL-4, IL-5, IL-6, IL-7, MIF, and TNFα), chemokines (BLC(CXCL13), Eotaxin (CCL11), I-TAC(CXCL11), IL-8(CXCL8), IP-10(CXCL10), MCP-1(CCL2), MIG(CXCL9), MIP-1a(CCL3), MIP-1b(CCL4)), growth factors (BDNF, G-CSF, GM-CSF, PDGF-BB, VEGF-A) and other factors (CD40L, Granzyme B) (Invitrogen, Carlsbad, CA). Differences in induction of proteins post stimulation were tested using unpaired t-tests with Welch’s correction. Dose-dependent responses were modeled based on g/kg/day ethanol consumed and tested for linear fit using regression analysis in Prism (GraphPad, San Diego CA). Raw data included in **Supp. Table 2**.

### RNA isolation and library preparation

Total RNA was isolated from PBMC or purified CD14+ monocytes using the mRNeasy kit (Qiagen, Valencia CA) following manufacturer instructions and quality assessed using Agilent 2100 Bioanalyzer. Libraries from PBMC RNA were generated using the TruSeq Stranded RNA LT kit (Illumina, San Diego, CA, USA). Libraries from purified CD14+ monocytes RNA were generated using the NEBnext Ultra II Directional RNA Library Prep Kit for Illumina (NEB, Ipswitch, MA, USA), which allows for lower input concentrations of RNA (10ng). For both library prep kits, rRNA depleted RNA was fragmented, converted to double-stranded cDNA and ligated to adapters. The roughly 300bp-long fragments were then amplified by PCR and selected by size exclusion. Libraries were multiplexed and following quality control for size, quality, and concentrations, were sequenced to an average depth of 20 million 100bp reads on the HiSeq 4000 platform.

### Bulk RNA-Seq data analysis

RNA-Seq reads were quality checked using FastQC (https://www.bioinformatics.babraham.ac.uk/projects/fastqc/), adapter and quality trimmed using TrimGalore(https://www.bioinformatics.babraham.ac.uk/projects/trim_galore/), retaining reads at least 35bp long. Reads were aligned to *Macaca mulatta* genome (Mmul_8.0.1) based on annotations available on ENSEMBL (Mmul_8.0.1.92) using TopHat (67) internally running Bowtie2 (68). Aligned reads were counted gene-wise using GenomicRanges (69), counting reads in a strand-specific manner. Genes with low read counts (average <5) and non-protein coding genes were filtered out before differential gene expression analyses. Read counts were normalized using RPKM method for generation of PCA and heatmaps. Raw counts were used to test for differentially expressed genes (DEG) using edgeR (70), defining DEG as ones with at least two-fold up or down regulation and an FDR controlled at 5%. Functional enrichment of gene expression changes in resting and LPS-stimulated cells was performed using Metascape (71) and DAVID (72). Networks of functional enrichment terms were generated using Metascape and visualized in Cytoscape (73). Transcription factors that regulate expression of DEG were predicted using the ChEA3 (74) tool using ENSEML ChIP database.

### 10X scRNA-Seq data analysis

Freshly thawed PBMC from control (n=3) and CHD (n=3) animals were stained with anti-CD14, HLA-DR antibodies and sorted for live CD14+/HLA-DR+ cells on a BD FACSAria Fusion. Sorted monocytes were pooled and resuspended at a concentration of 1200 cells/ul and loaded into the 10X Chromium gem aiming for an estimated 10,000 cells per sample. cDNA amplification and library preparation (10X v3 chemistry) were performed according to manufacturer protocol and sequenced on a NovaSeq S4 (Illumina) to a depth of >50,000 reads/cell.

Sequencing reads were aligned to the Mmul_8.0.1 reference genome using cellranger v3.1 (75) (10X Genomics). Quality control steps were performed prior to downstream analysis with *Seurat*, filtering out cells with fewer than 200 unique features and cells with greater than 20% mitochondrial content. Control and CHD datasets were integrated in Seurat (76) using the *IntegrateData* function. Data normalization and variance stabilization were performed, correcting for differential effects of mitochondrial and cell cycle gene expression levels. Clustering was performed using the first 20 principal components. Small clusters with an over-representation of B and T cell gene expression were removed for downstream analysis. Clusters were visualized using uniform manifold approximation and projection (UMAP) and further characterized into distinct monocyte subsets using the *FindMarkers* function (**Supp. Table 3**).

### Pseudo-temporal analysis

Pseudotime trajectory monocytes was reconstructed using Slingshot (77). The UMAP dimensional reduction performed in Seurat was used as the input for Slingshot. For calculation of the lineages and pseudotime, the most abundant classical monocyte cluster, MS1, was set as the root state.

### Differential expression analyses

Differential expression analysis (CHD to Control) was performed using *DESeq* under default settings in *Seurat*. Only statistically significant genes (Fold change cutoff ≥ 1.2; adjusted p-value ≤ 0.05) were included in downstream analysis.

### Module Scoring and functional enrichment

For gene scoring analysis, we compared gene signatures and pathways from KEGG (https://www.genome.jp/kegg/pathway.html) (**Supp. Table 4**) in the monocytes using Seurat’s *AddModuleScore* function. Over representative gene ontologies were identified using 1-way, 2-way or 4-way enrichment of differential signatures using Metascape (71). All plots were generated using *ggplot2* and *Seurat*.

### ATAC-Seq library preparation

Following the Omni-ATAC protocol, 2×10^4^ purified monocytes were lysed in lysis buffer (10mM Tris-HCl (pH 7.4), 10mM NaCl, 3mM MgCl_2_, 10% Np-40, 10% Tween, and 1% Digitonin) on ice for 3 minutes (42). Immediately after lysis, nuclei were spun at 500 g for 10 minutes at 4C to remove supernatant. Nuclei were then incubated with Tn5 transposase for 30 minutes at 37C. Tagmented DNA was purified using AMPure XP beads (Beckman Coulter, Brea, CA) and PCR was performed to amplify the library under the following conditions: 72C for 5 min; 98 for 30s; 5 cycles of 98C for 10s, 63C for 30s, and 72C for 1min; hold at 4C. Libraries were then purified with warm AMPure XP beads and quantified on a Bioanalyzer (Agilent Technologies, Santa Clara CA). Libraries were multiplexed and sequenced to a depth of 50 million 100bp paired reads on a NextSeq (Illumina).

### ATAC-Seq data analysis

Paired ended reads from sequencing were quality checked using FastQC and trimmed to a quality threshold of 20 and minimum read length 50. Trimmed reads were aligned to the Macaca Mulatta genome (Mmul_8.0.1) using Bowtie2 (-X 2000 -k 1 --very-sensitive --no-discordant --no-mixed). Reads aligning to mitochondrial genome were removed using Samtools and PCR duplicate artifacts were removed using Picard. Samples from each group were concatenated to achieve greater than 5×10^6^ non-duplicate, non-mitochondrial reads per group.

Accessible chromatin peaks were called using Homer’s *findPeaks* function (78) (FDR<0.05) and differential peak analysis was performed using Homer’s *getDifferentialPeaks* function (P < 0.05). Genomic annotation of open chromatin regions in monocytes and differentially accessible regions (DAR) with CHD was assigned using ChIPSeeker (79). Promoters were defined as −1000bp to +100bp around the transcriptional start site (TSS). Functional enrichment of open promoter regions was performed using Metascape (http://metascape.org).

Due to the lack of available macaque annotation databases, distal intergenic regions from the macaque assembly were converted to the human genome (hg19) coordinates using the UCSC liftOver tool. *Cis*-Regulatory roles of these putative enhancer regions were identified using GREAT (http://great.stanford.edu/public/html/). Transcription factor motif analysis was performed using Homer’s *findMotifs* function with default parameters. Promoter regions for every annotated macaque gene were defined in ChIPSeeker as -1000bp to +100bp relative to the TSS. A counts matrix was generated for these regions using *featureCounts (80)*, where pooled bam files for each group were normalized to total numbers of mapped reads.

### Phagocytosis Assay

500,000 freshly thawed total BAL cells were resuspended in RP10 media supplemented with 100ng/mL LPS and incubated for 4 hours at 37C with 5% CO_2_. 50uL of pHrodo Red S.aureus BioParticles (Thermo Fisher Scientific, Waltham, MA) were added to the cells and they were incubated for an additional 2 hours in the incubator. The cells were washed and stained with anti-CD206 antibody and acquired with an Attune NxT Flow Cytometer (ThermoFisher Scientific, Waltham, MA) and further analyzed using FlowJo software (Ashland, OR).

### Statistical Analysis

All statistical analyses were conducted in Prism 7(GraphPad). Data sets were first tested for normality. Two group comparisons were carried out an unpaired t-test with Welch’s correction. Differences between 4 groups were tested using one-way ANOVA (α=0.05) followed by Holm Sidak’s multiple comparisons tests. Error bars for all graphs are defined as ± SEM. Linear regression analysis compared significant shifts in curve over horizontal line, with spearman correlation coefficient reported. Statistical significance of functional enrichment was defined using hypergeometric tests. P-values less than or equal to 0.05 were considered statistically significant. Values between 0.05 and 0.1 are reported as trending patterns.

## Author Contributions

S.A.L., K.A.G., and I.M. conceived and designed the experiments. S.A.L., S.S., and B.D. performed the experiments. S.A.L. and B.D. analyzed the data. S.A.L. and I.M. wrote the paper. All authors have read and approved the final draft of the manuscript.

## Acknowledgements

We are grateful to the members of the Grant laboratory for expert animal care and sample procurement. We thank Dr. Jennifer Atwood for assistance with sorting in the flow cytometry core at the Institute for Immunology, UCI. We thank Dr. Melanie Oakes from UCI Genomics and High-Throughput Facility for assistance with 10X library preparation and sequencing.

## Funding

This study was supported by NIH 1R01AA028735-01 (Messaoudi), 5U01AA013510-20 (Grant), and 2R24AA019431-11 (Grant). S.A.L is supported by NIH 1F31A028704-01. The content is solely the responsibility of the authors and does not necessarily represent the official views of the NIH.

## Competing interests

No competing interests reported.

## Data availability

The datasets supporting the conclusions of this article are available on NCBI’s Sequence Read Archive (SRA# PRJNA723053).

## Notes

### Competing Interest Statement

The authors have declared no competing interest.

