## Supplemental Figures for "Transcriptional, epigenetic, and functional reprogramming of blood monocytes in non-human primates following chronic alcohol drinking"

**SUPPLEMENTARY FIGURES**:

Supplemental Figure 1

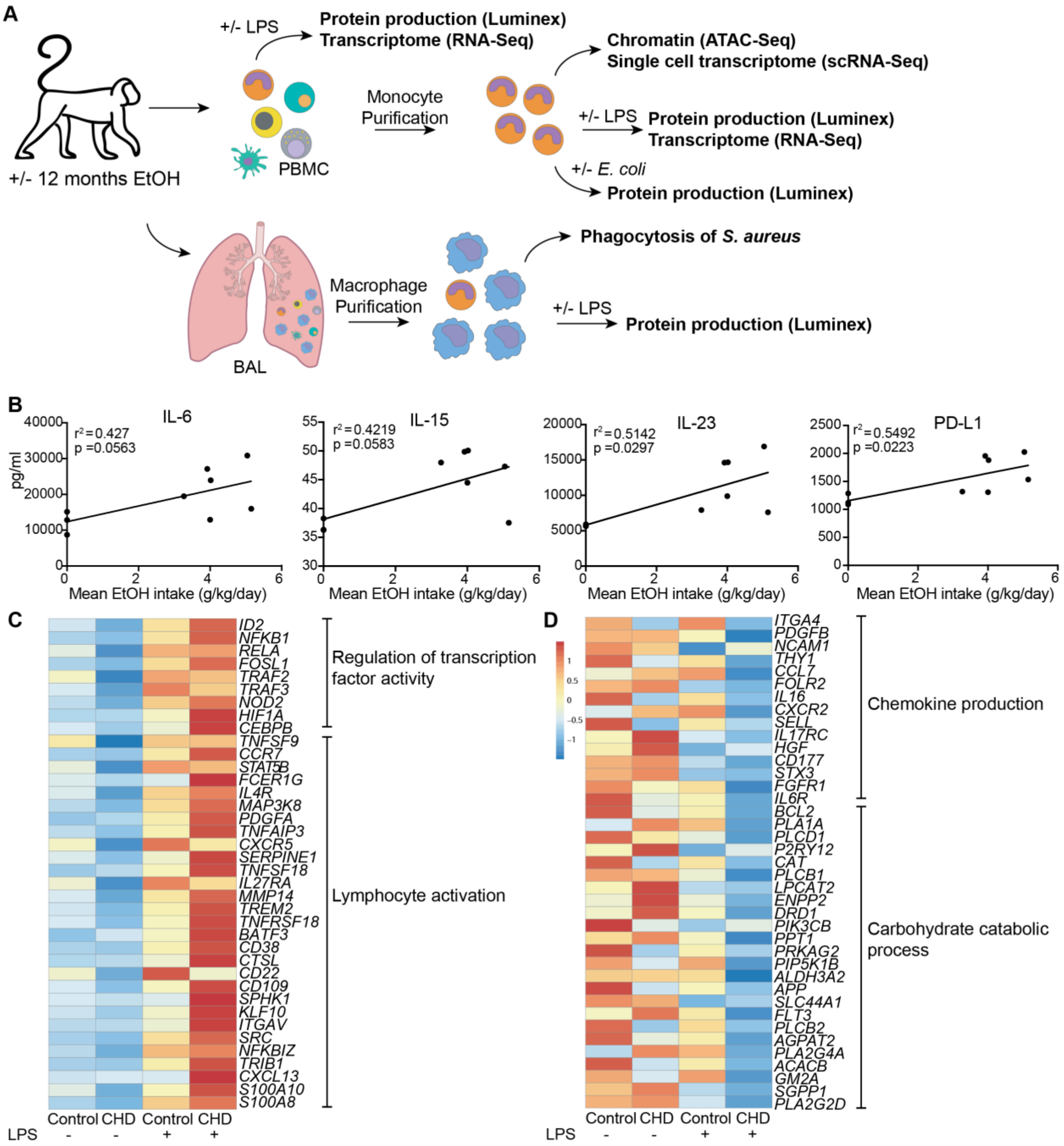


**Supp. Figure 1**: A) Experimental Design figure for this study. B) Scatter plots showing Spearman correlation between average EtOH dose (grams EtOH/kg body weight/day) and concentration (pg/ml) of the secreted factors IL-6, IL15, IL-23, PD-L1. C) Heatmap of genes related to *Regulation of transcription factor activity* and *Lymphocyte activation* upregulated in CHD PBMC with LPS. D)

Heatmap of genes involved in *Chemokine production* and *Carbohydrate catabolic process* pathways downregulated in CHD PBMC with LPS.

Supp. Figure 2

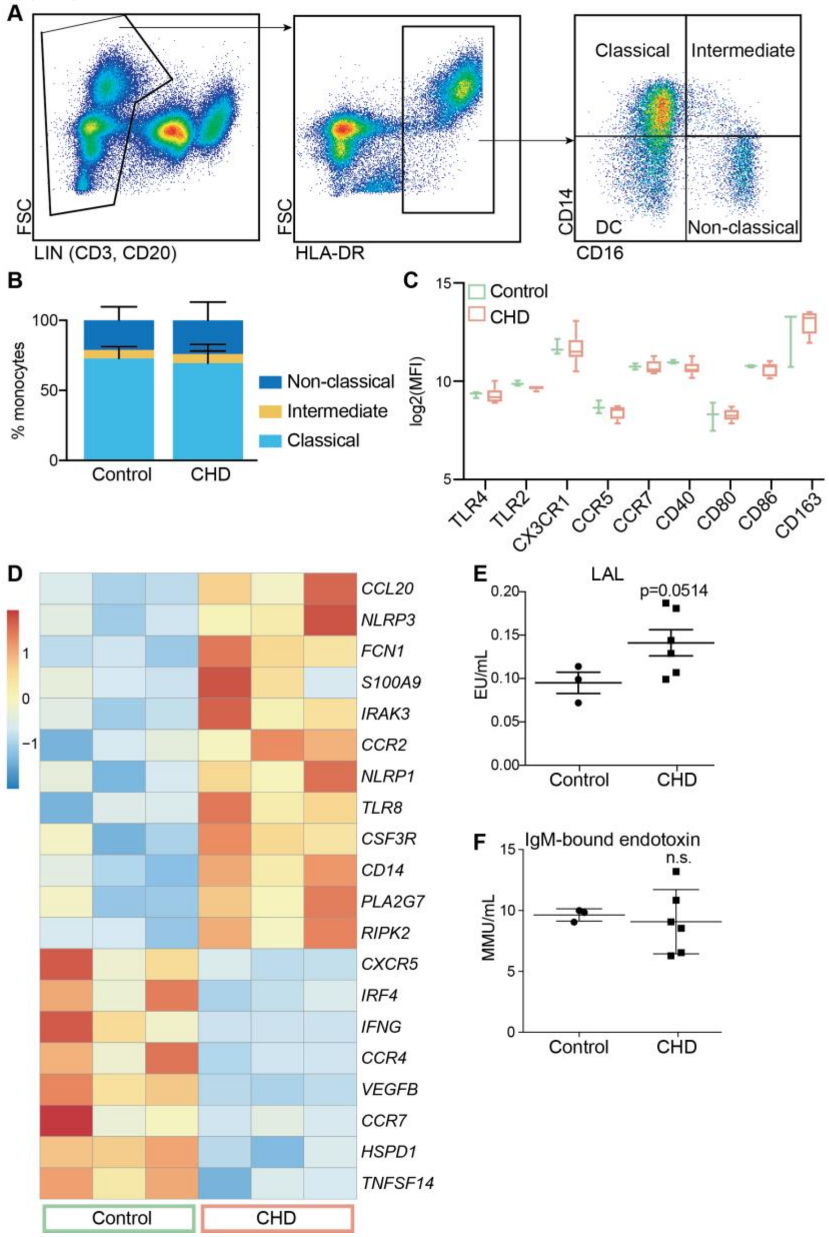


**Supp. Figure 2**: A) Gating strategy used to identify the three populations of monocytes in the blood. B) Relative percentages of the three monocyte populations in the controls and CHD macaques. C) $Log_2$ median fluorescence intensities (MFI) of monocyte cell surface markers across the controls and CHD macaques. D) Heatmap of normalized expression of differentially expressed genes between resting control and CHD monocytes. E,F) LAL (E) and IgM-bound endotoxin (F) levels measured from plasma by ELISA. T-test with Welch's correction was used to measure significance.

Supplemental Figure 3

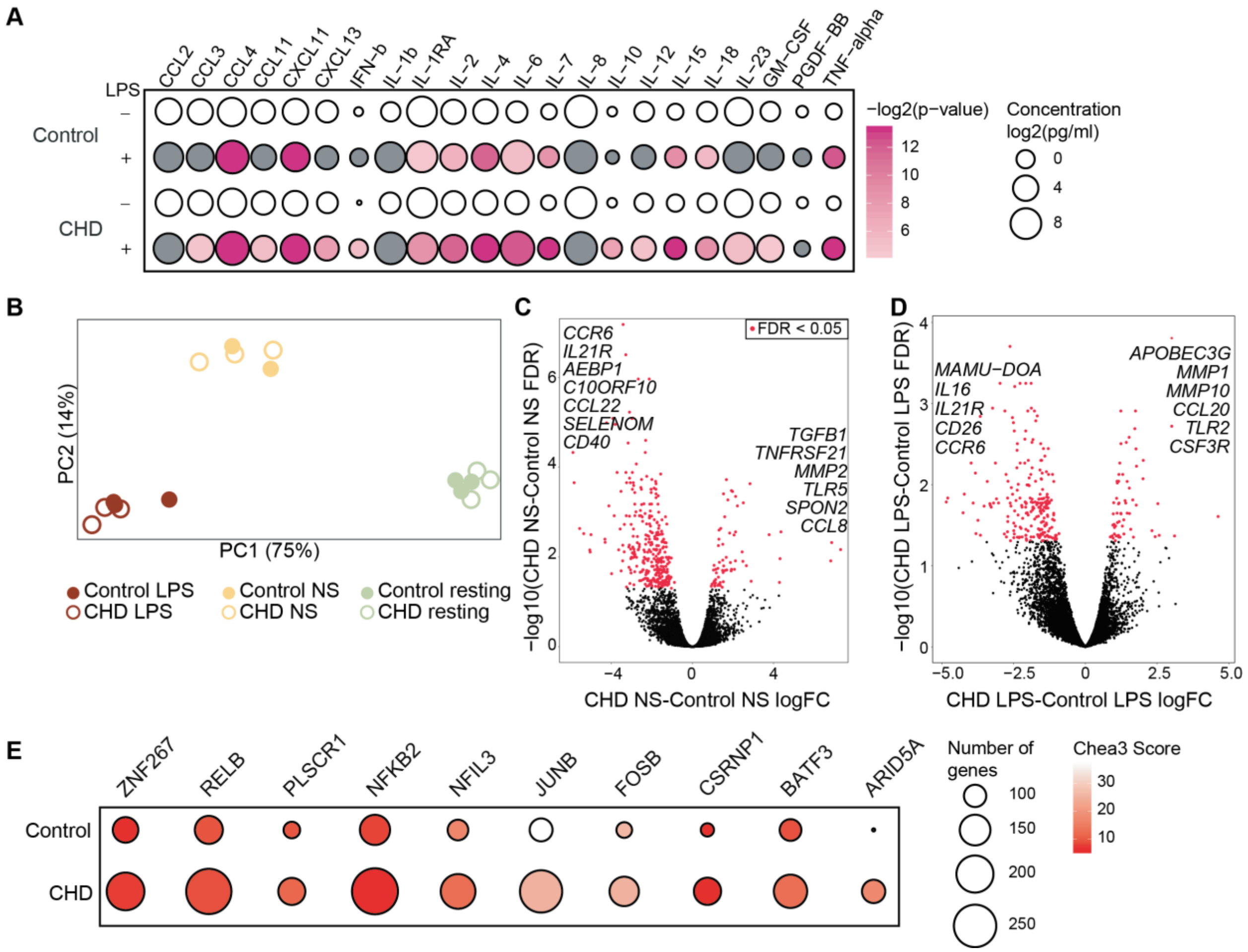


**Supp. Figure 3**: A) Bubble plot representing immune factor production (pg/ml) in the presence or absence of LPS stimulation of monocytes from control and CHD animals. The size of each circle represents the indicates the $log_2$ mean concentration of the indicated secreted factor and the color denotes the -$log_2$ transformed p value with the darkest pink representing the most significant value. The p-values were calculated between the unstimulated and stimulated conditions for each group using One-way ANOVA and a p-value cut-off of 0.05 was set. White circles indicate non-significant p-value. B) Principal component analysis (PCA) of CHD and control monocytes under resting, unstimulated (6 hours of media) and LPS stimulation (6 hours) conditions. C,D) Volcano plot representing up- and downregulated DEG with CHD in non-stimulated (C) and LPS stimulated (D) monocytes. Red = significant with an FDR ≤ 0.05 and fold change ≥ 2. E) Bubble plot representing transcription factors predicted to regulate LPS-responsive DEGs. The size of the dot represents the number of genes and the color represents Chea3 score where lower number is more significant. Analysis was performed using the Chea3 web browser.

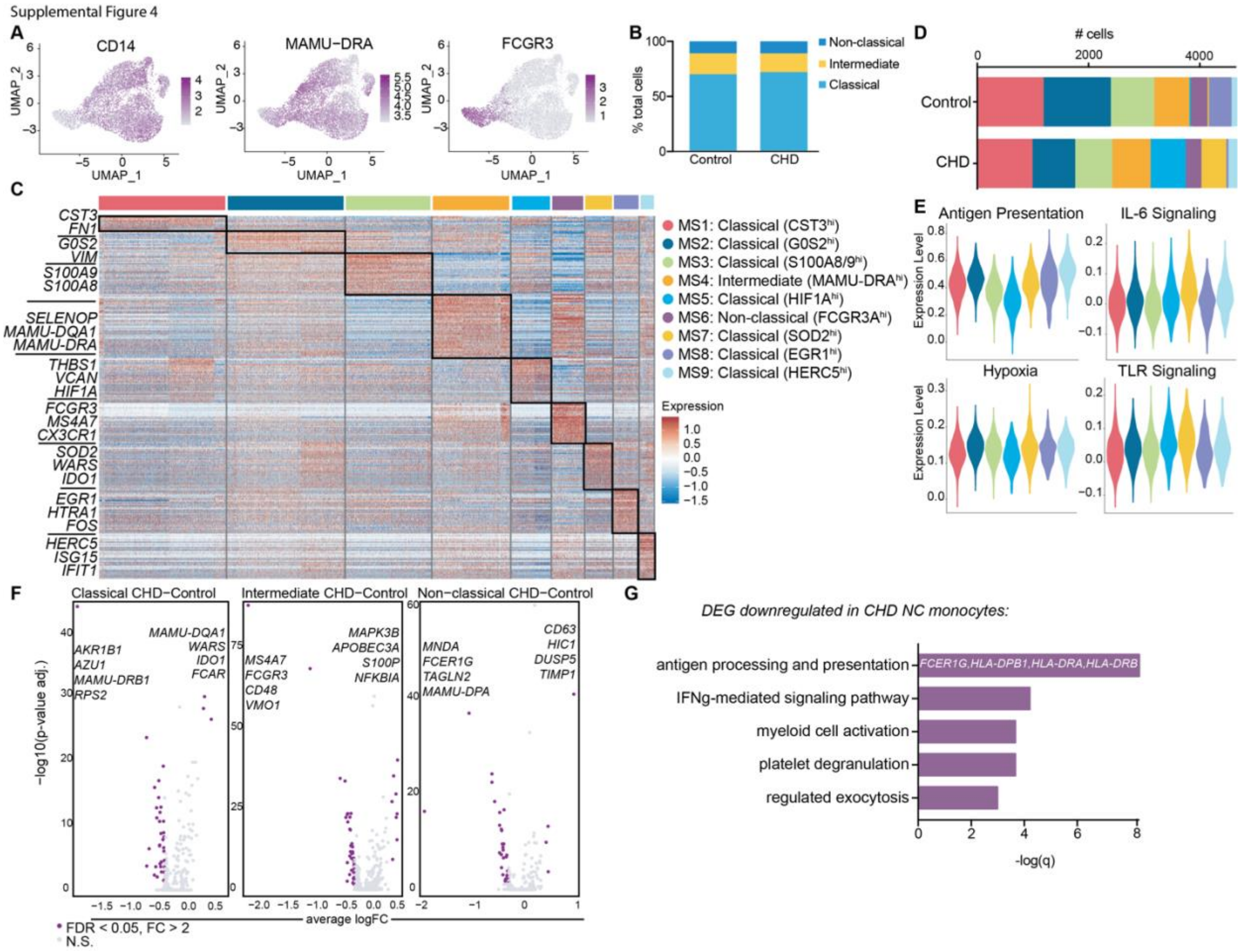


**Supp. Figure 4**: A) Feature plots showing relative gene expression of canonical monocyte markers *CD14*, *MAMU-DRA*, and *FCGR3* in all monocytes. B) Stacked bar graph depicting relative abundance of non-classical, intermediate, and classical monocytes in control and CHD monocytes. C) Heatmap showing relative gene expression of the representative genes used for clustering and subset identification. D) Stacked bar graph depicting abundance of cells clustered in each monocyte subset in control and CHD groups. E) Violin plots comparing antigen presentation, IL-6 signaling, hypoxia, and TLR signaling module scores with the classical monocyte subsets. F) Volcano plots of the up-and downregulated genes comparing CHD to control non-classical, intermediate, and classical monocytes. The purple color indicates significant DEG where FDR ≤ 0.05 and fold-change ≥ 2. G) Bar graph showing functional enrichment pathways of DEG downregulated with CHD in the non-classical monocyte cluster (MS6), defined by q-value.

Supplementary Figure 5

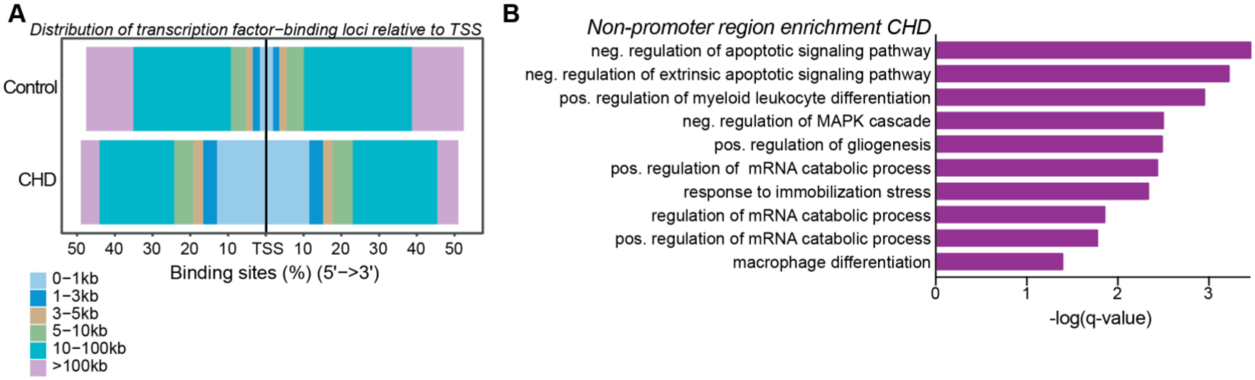


**Supp. Figure 5**: A) Distribution of the open chromatin regions in the CHD and control monocytes relative to the transcription start site (TSS). B) Top GO Biological process enrichment of the genes predicted to be cis-regulated by non-promoter regions more open with CHD. Genomic regions were lifted over from rhesus macaque to homo sapien genome (UCSC) and further enriched using the GREAT.

**SUPPLEMENTARY TABLES**:

**Supp. Table 1**: Animals and EtOH g/kg values

**Supp. Table 2**: Immune mediator production by PBMC, purified monocytes, and AM following LPS or *E.coli* stimulation.

**Supp. Table 3**: Genes associated with each MS cluster

**Supp. Table 4**: Module Scoring genes
