## Supplementary material for "Transcriptional, epigenetic, and functional reprogramming of blood monocytes in non-human primates following chronic alcohol drinking": Supp. Table 1

| **Animal ID** | **Sex** | **Mean daily ethanol intake (g/kg/day)** | **Standard Deviation of mean daily intake** | **Blood**  **Ethanol**  **Content**  **(Avg mg%)** | **Drinking status** |
| --- | --- | --- | --- | --- | --- |
| C1 | F | 0 | - | 0 | Control |
| C2 | F | 0 | - | 0 | Control |
| C3 | F | 0 | - | 0 | Control |
| C4 | M | 0 | - | 0 | Control |
| C5 | M | 0 | - | 0 | Control |
| C6 | M | 0 | - | 0 | Control |
| C7 | M | 0 | - | 0 | Control |
| CHD1 | F | 3.8 | 1.0 | 41 | EtOH |
| CHD2 | F | 4.1 | 1.0 | 46 | EtOH |
| CHD3 | F | 4.2 | 1.1 | 57 | EtOH |
| CHD4 | F | 4.2 | 1.3 | 65 | EtOH |
| CHD5 | F | 5.3 | 1.4 | 81 | EtOH |
| CHD6 | F | 5.6 | 1.2 | 103 | EtOH |
| CHD7 | M | 2.9 | 1.3 | 74 | EtOH |
| CHD8 | M | 3.2 | 0.9 | 79 | EtOH |
| CHD9 | M | 3.0 | 0.6 | 79 | EtOH |
| CHD10 | M | 3.3 | 1.0 | 98 | EtOH |

**Table 1: Summary of samples used in this study.** Mean daily ethanol (EtOH) intake reflects the average dose consumed during the period of 12-month self-administration period.
